# Whole-Mount Optical Clearing of Rabbit Tenuissimus Muscle for Assessment of Muscle Spindle Morphology

**DOI:** 10.64898/2026.04.28.721525

**Authors:** Emily J Reedich, Brendan Moline, Oluwatobi Opesade, Cassandra Kramer, Jess Glennon, Eden Fraatz, Katharina Quinlan, Marin Manuel

## Abstract

Proprioception and reflexive control of muscle tone depend on the activity of muscle spindles, specialized sensory receptors embedded deep within skeletal muscle that detect changes in muscle length. Their location and complex three-dimensional architecture have historically limited morphological analysis to techniques such as silver-impregnation, muscle teasing, or serial sectioning followed by volumetric reconstruction. Here, we describe a workflow for three-dimensional, in situ visualization of muscle spindles in the rabbit tenuissimus muscle, a preparation uniquely enriched in spindles and well suited for whole-mount imaging. The protocol combines fluorescent labeling of spindle sensory and motor innervation, including intrafusal γ neuromuscular junctions labeled with α-bungarotoxin, with immunolabeling and solvent-based optical clearing. Optically cleared tenuissimus muscles were compatible with both whole-mount confocal and light-sheet microscopy, enabling volumetric imaging of complete spindle structures and detailed visualization of Ia annulospiral endings at the spindle equator. This approach provides access to spindle morphology and connectivity at multiple spatial scales while avoiding physical sectioning and reconstruction. By enabling reproducible three-dimensional imaging of intact muscle spindles, this workflow offers a practical platform for studying spindle structure and plasticity in health and disease.

## 1 Introduction

Muscle spindles are specialized sensory receptors embedded in skeletal muscle that detect changes in muscle length and relay that information to the central nervous system via afferent fibers (Ia and II). They thereby contribute to proprioception and reflex control of muscle tone, forming the afferent limb of the stretch (myotatic) reflex loop (Pierrot-Deseilligny and Burke, 2012). The spindle is composed of intrafusal muscle fibers (two types of nuclear bag and one type of nuclear chain fibers), with contractile polar regions and a noncontractile equatorial region where the sensory endings terminate (Banks, 2023, 2015). The sensitivity of the spindle can be modulated by γ motoneuron (fusimotor) innervation at γ neuromuscular junctions: activation of γ fibers (dynamic or static) changes the tension of intrafusal fibers and thus alters the responsiveness of the sensory afferents to stretch (Banks, 2015; Bergenheim et al., 1995; Bishop et al., 1968; Matthews, 1958). Because spindles provide essential feedback for motor control, their structural integrity and proper innervation are critical for normal movement, reflex responses, and adaptation (e.g., motor learning, postural control) (Bartels and Harrison, 2025; Santuz and Akay, 2023).

The tenuissimus muscle in certain species (e.g. cat, rabbit, hamster) has historically been used as a model preparation in spindle physiology and fusimotor studies. The thin, elongated architecture and relative simplicity of this muscle is ideal to study muscle spindles. Their particular arrangement in the muscle allows spindles and sensory and motor axons to be traced more easily than in denser muscles (Butler, 1980). Because the tenuissimus is a relatively thin muscle, light scattering and penetration issues are more manageable in whole-mount imaging. With the advent of tissue clearing techniques (solvent-based, aqueous-based, and hydrogel-based; (Ueda et al., 2020)), this approach can be further refined to offer unprecedented access to the structure of muscle spindles in healthy and disease models.

One hurdle to muscle spindle visualization in optically cleared skeletal muscle is the reported incompatibility between the primary reagent for visualizing γ neuromuscular junctions [fluorescent dye-conjugated α-bungarotoxin (BTX)] and key chemicals involved in clearing (Zhan et al., 2023). No previous study has combined α-BTX labeling of γ endplates with iDISCO and solvent-based clearing in skeletal muscle. In this study, we adapt a tissue clearing + immunofluorescence pipeline to the tenuissimus muscle, allowing the visualization of the entire architecture of muscle spindles in three dimensions. Given the central role of spindles in motor control, our approach provides a novel way to study muscle spindle morphology, connectivity, and plasticity in health and disease, *in situ*.

## 2 Methods

### 2.1 Muscle dissection

#### Ethics statement

All experimental procedures involving animals were approved by the University of Rhode Island’s Institutional Animal Care and Use Committee and performed in accordance with the National Institutes of Health *Guide for the Care and Use of Laboratory Animals*.

#### Animal housing and husbandry

Neonatal New Zealand White rabbit kits were born to dams bred in-house in our colony that is derived from Charles River Laboratories, Inc. (Wilmington, MA). Rabbits were housed in a controlled vivarium with a 12:12 h light-dark cycle and had access to food and water *ad libitum*.

#### Isolation of the tenuissimus muscle

Rabbit kits were euthanized in the first two postnatal weeks by intraperitoneal injection of EUTHASOL Euthanasia Solution (pentobarbital sodium and phenytoin sodium; Patterson Veterinary, cat. # 07-805-9296) followed by decapitation. The fur over the hindlimb was wetted with ethanol and removed to expose the underlying musculature. The entire hindlimb was dissected from the rest of the carcass and pinned to a Sylgard-lined petri dish in a lateral recumbent orientation with the knee joint at a 90° angle (Figure 1A). The hindlimb was submerged in chilled phosphate buffered saline (1x PBS).

**Figure 1:**
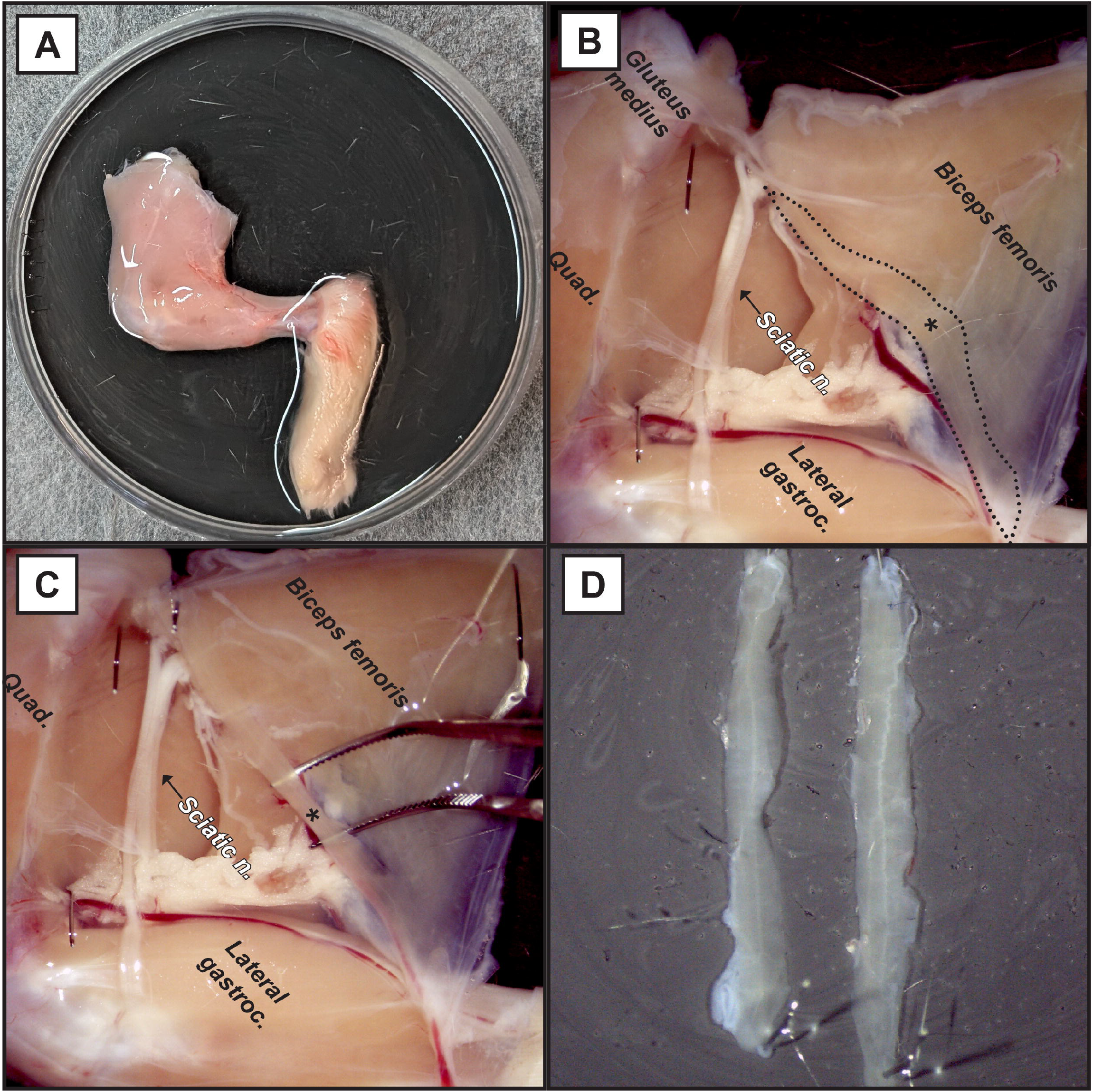
Dissection of the tenuissimus muscle. **(A)** The neonatal rabbit hindlimb is pinned in a lateral recumbent position to a Sylgard-lined petri dish with the knee joint at a 90 ° angle. **(B)** The tenuissimus (*) is visible upon medial reflection of the biceps femoris, with which it shares a tendon of insertion along the tibia. **(C)** The tenuissimus (*) is isolated by blunt dissection from the biceps femoris and proximal-to-distal resection. **(D)** Two examples of the isolated tenuissimus; a characteristic neurovascular bundle runs along its midline. Abbreviations: gastroc., gastrocnemius; n., nerve; quad., quadriceps femoris.

An incision was made by the patella between the fascia lata and the biceps femoris muscle. The incision was extended medially to the caudal fascia, separating the tensor fasciae latae and gluteus medius muscles from the gluteus maximus. The original incision by the patella was extended through the biceps femoris tendon of insertion and along the tibia distally to the Achilles tendon. The transected biceps femoris tendon of insertion was held with forceps and reflected towards the midline to expose the tenuissimus muscle (Figure 1B), which originates at the transverse process of the second caudal vertebra and shares its transected tendon of insertion with the biceps femoris (Chin, 1957; Patten and Ovalle, 1992; Walker, 1980). The tenuissimus was gently blunt dissected along the biceps femoris (Figure 1C); next, it was held by its tendon of origin and isolated by resecting it distally to its transected tendon of insertion.

### 2.2 Optical Clearing

#### Labeling intrafusal fiber motor end plates with α-BTX and tissue fixation

The isolated tenuissimus was immediately transferred to a microfuge tube (5 mL) containing α-bungarotoxin (α-BTX)-Alexa Fluor 647 conjugate (Life Technologies, cat. # B35450) diluted 1:250 in 1x PTwH, a solution made of 1x PBS with Tween-20 (0.2%) and heparin (10 μg/mL; Sigma cat. #H3393) and incubated on a shaker plate at room temperature in the dark for 1 h to fluorescently label motor end plates. All proceeding incubations were also performed in the dark. The muscle was washed once quickly then 4 x 5 minutes in chilled 1x PBS on a shaker plate at room temperature, and transferred to a Sylgard-lined petri dish. Insect pins were used to hold the tenuissimus in a linear position in the dish (Figure 1D); care was taken to ensure the muscle was not stretched. The muscle was then immersion-fixed in 4% paraformaldehyde (in 1x PBS; pH 7.4) at 4 °C overnight. The next day, the tissue was washed 3 x 10 minutes in 1x PBS and stored in 1x PBS with sodium azide (0.05%) until immunolabeling. Of note, we tested variations in α-BTX application, and found that increased concentration (1:100), lowered temperature (4 °C for 2 h), and *in vitro* incubation (room temperature for 4 h) in carbogenated physiological saline (formulated as in (Reid et al., 1999)) all yielded similar fluorescence intensity of α-BTX labeling.

#### Immunolabeling of annulospiral endings

A methanol gradient was used to permeabilize tenuissimus muscles for immunolabeling. We adapted our methanol pre-treatment protocol from (Renier et al., 2014), choosing to omit the hydrogen peroxide-based tissue bleaching step in order to preserve α-BTX-Alexa Fluor 647 fluorescence (Renier et al., 2014). The samples were incubated (all volumes: 1 mL) on a shaker plate at room temperature, for 1 h each, in: 50% methanol (diluted in 1x PBS), 80% methanol (diluted in 1x PBS); then 2 x 1 h in 100% methanol. The sample was next incubated on a shaker plate at 4 °C overnight in 20% dimethylsulfoxide (DMSO; in methanol). The next day, the sample was incubated on a shaker plate at room temperature for 1 h in 50% methanol (diluted in 1x PBS), followed by 2 x 1 h in 1x PBS then 2 x 1 h in 0.2% Triton X-100.

To visualize the axons, we performed immunolabeling of neurofilament and synaptic vesicle glycoprotein 2, which labels motor axons, their terminals, and sensory axons including Ia annulospiral endings. We adapted the protocol from (Renier et al., 2014). Immediately after pretreatment, samples were incubated (all volumes: 1 mL) in 0.2% Triton X-100, 20% DMSO, 0.3 M glycine (diluted in 1x PBS) inside an incubator-shaker at 37 °C overnight. The samples were blocked in 0.2% Triton X-100, 10% DMSO, 6% goat serum (diluted in 1x PBS) inside an incubator-shaker at 37 °C for one day. The samples were washed on a shaker plate at room temperature for 2 x 1 h in PTwH. The samples were incubated in primary antibody cocktail (diluted in 5% DMSO, 3% goat serum in PTwH) inside an incubator-shaker at 37 °C for two days. Primary antibody cocktail consisted of mouse anti-synaptic vesicle 2 (DSHB, cat. # SV2-s; diluted 1:20) and mouse anti-neurofilament (DSHB, cat. # 2H3-s; diluted 1:100). The samples were washed on a shaker plate at room temperature for one day in PTwH (the buffer was replaced at 1 h, 2 h, and 4 h marks). Next, the samples were incubated in secondary antibody solution (diluted in 3% goat serum in PTwH) inside an incubator-shaker at 37 °C for one day. Secondary antibody solution was composed of Alexa Fluor 555 AffiniPure Goat Anti-Mouse IgG (H+L) (Jackson ImmunoResearch; cat. #115-565-146; diluted 1:500). Lastly, the samples were washed on a shaker plate at room temperature for one day in PTwH (the buffer was replaced at 1 h, 2 h, and 4 h marks).

#### Optical clearing

We adapted FDISCO (Qi et al., 2019a) for solvent-based optical clearing of fluorescently labeled tenuissimus muscles. Samples were delipidated and dehydrated by incubation on a shaker plate at 4°C, for 1.5 h each (all volumes: 1 mL), in: 50% tetrahydrofuran (THF), 0.01% triethylamine in ultrapure water; 70% THF, 0.01% triethylamine in ultrapure water; 80% THF, 0.01% triethylamine in ultrapure water. The samples were then incubated in 100% THF, 0.01% triethylamine on a shaker plate at 4 °C for 2 x 2 h. For refractive index matching, the samples were incubated (volume: 5 mL) in 100% dibenzyl ether (DBE) at 4 °C for 1 h; this step rendered the samples transparent. The samples were stored in a fresh aliquot of 100% DBE in the dark at 4 °C until microscopy. We note that our samples were also amenable to room temperature storage and imaging in the non-toxic alternative mounting medium, ethyl cinnamate (100%).

### 2.3 Confocal microscopy

For confocal microscopy, the samples were mounted onto microscope slides (Globe Scientific, Inc. #1380-20) using the solvent used for refraction index matching (100% DBE or 100% ethyl cinnamate) as the mounting medium. For thicker samples, a thin barrier of vacuum grease can be applied around the coverlip (#1.5; VWR # 48393-241) to seal in the mounting medium.

Widefield images were acquired on a Nikon ECLIPSE Ti2-E inverted microscope with a motorized stage and a Sola light engine (Lumencor). Observations were performed using a Nikon CFI Plan Apochromat Lambda D 10×/0.45NA air objective, a DsRed/TRITC/Cy3 Filter Set (Nikon #96394, excitation: 554/23, dichroic: 573, emission: 609/54) for Alexa Fluor 555, and a Cy5 filter set (Nikon #96232, excitation:618/50, dichroic: 652, emission: 698/70) for Alexa Fluor 647. Images were acquired using a Zyla 4.2 sCMOS camera (Andor; 12bit, gain 4, rolling shutter, no binning) and NIS-Elements AR v.6.10.01 (Nikon).

Confocal Z-stacks were acquired on a Nikon ECLIPSE Ti2-E inverted microscope with a motorized stage and a LUN4 laser engine (405/488/561/640nm, 15mW), equipped with a Nikon C2+ galvo scanning head, and two C2-DUVB GaAsP detector units. Images were acquired with a CFI Plan Apochromat Lambda D 20×/0.8 air objective, or a CFI Plan Apochromat Lambda D 40×/0.95 air objective, optionally with a 1.5× zoom lens inserted in the light path, and NIS-Elements AR v.6.10.01 (Nikon). Details about acquisition parameters are provided in the legends of the relevant figures.

Lightsheet volumetric images were acquired on either a MegaSPIM (LifeCanvas Technology, Cambridge, MA) or a ZEISS Lightsheet 7 light□sheet fluorescence microscope (Carl Zeiss, Germany). The MegaSPIM illumination and detection are performed using long□working□distance optics positioned at a 45° angle relative to the sample plane. Excitation was provided using a 561□nm laser line. Fluorescence was detected using a 22×/0.7□NA multi□immersion dipping objective. Emitted fluorescence was captured using a Hamamatsu ORCA□Fusion sCMOS camera (2304□×□2304□pixels). Raw image stacks were computationally processed using the MegaSPIM post□processing pipeline, including deskewing and stitching, to generate isotropic three□dimensional image volumes suitable for downstream visualization and quantitative analysis. The movie was generated using Arivis Pro v.4.2.1.

The Lightsheet 7 uses two illumination objectives and perpendicular detection optics. Illumination was provided via fused dual light-sheets, with pivot scanning enabled. Images were detected using either a 5×/0.16NA or a 20×/1.0NA Plan□Neofluar objective, coupled to a PCO.Edge 4.2 sCMOS camera (2048□×□2048□px). Laser excitation was performed at 561, and 635□nm; signals were separated using an LBF 405/488/561/640 filter cube; images were acquired in two sequential tracks in continuous Z□stack mode with 10□% tile overlap, pivot scan enabled, and track switching via Z□stack multitrack acquisition, implemented through Zen Black 3.0 software.

## 3 Results

### Solvent-based optical clearing enables visualization of fluorescently labeled sensory and motor innervation of muscle spindles

The tenuissimus is a long, thin, fusiform muscle that spans across both the knee and the hip in many species. It is thought to play an important role in proprioceptive sensory feedback for the hindlimb (Patten and Ovalle, 1992). In this study, we developed an immunofluorescence and optical clearing workflow for three-dimensional visualization of muscle spindles in the rabbit tenuissimus (Figure 2). We labeled spindle poles by staining γ-neuromuscular junctions with fluorescent α-BTX and chemically cross-linked this reagent in place by immersion fixation in paraformaldehyde. Then, we adapted the immunofluorescence framework of iDISCO (Renier et al., 2014) to fluorescently label neurofilament and synaptic vesicle glycoprotein 2 for visualization of spindle sensory and motor innervation. Finally, we adapted the solvent-based optical clearing pipeline of FDISCO (Qi et al., 2019a) to achieve dehydration, delipidation, and refractive index matching of the fluorescently labeled tenuissimus. Our optical clearing protocol resulted in robust tissue transparency and slight shrinkage (18.8 ± 4.7% (n=3), Figure 3). As such, our workflow enables volume imaging of the optically cleared tenuissimus (Supplemental Movie 1).

**Figure 2:**
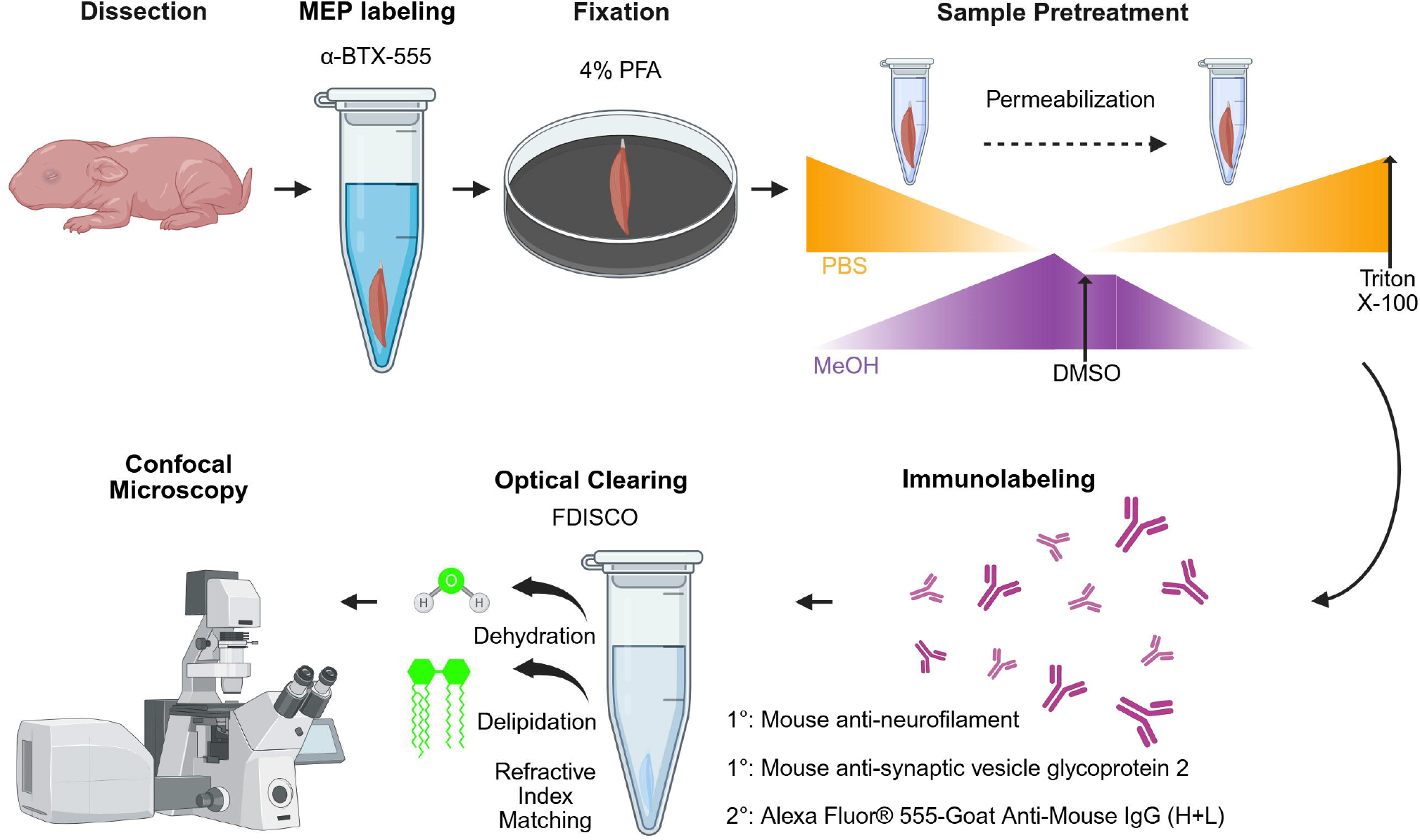
Immunolabeling and optical clearing workflow for three-dimensional visualization of muscle spindles in the tenuissimus. The tenuissimus muscle is dissected from the neonatal rabbit and incubated in α-BTX-Alexa Fluor 647 to label γ-neuromuscular junctions at spindle poles; then, the sample is pinned flat and chemically fixed by immersion in paraformaldehyde. After fixation, the tenuissimus is permeabilized for immunofluorescence by methanol pretreatment. Antibodies against neurofilament and synaptic vesicle glycoprotein 2 are used to immunolabel spindle sensory and motor innervation. The solvent-based optical clearing protocol FDISCO (Qi et al., 2019a) is implemented to achieve tissue dehydration, delipidation, and refractive index matching of the fluorescently labeled tenuissimus. Muscle spindles in the optically cleared tenuissimus are visualized and volume-imaged using confocal or light-sheet microscopy.

**Figure 3:**
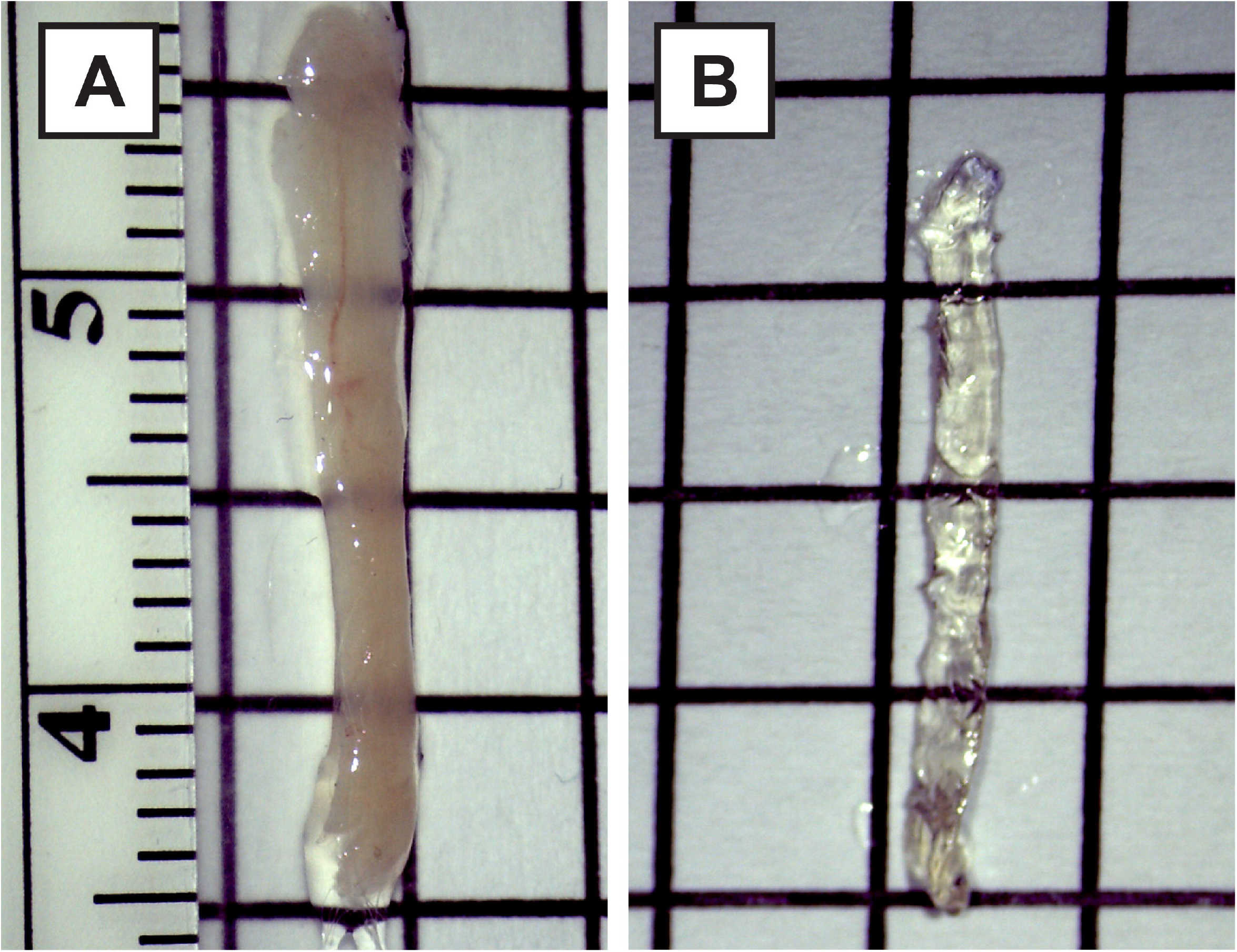
The isolated neonatal rabbit tenuissimus muscle. **(A)** Before and **(B)** after solvent-based optical clearing of a 12-day-old rabbit tenuissimus by FDISCO. In both images, each square is 5×5 mm.

Using this technique, we were able to visualize spindle structures in their entirety. We observed a high density of spindles near the midline of the muscle, parallel to the central neurovascular bundle (Figure 4A). The equatorial region of the spindles is easily identifiable by the neurofilament labeling of the annulospiral endings (Figure 4B–C). Spindle poles were identified by labeling γ-neuromuscular junctions with fluorescent α-BTX. Although this reagent is incompatible with common optical clearing methods (Milgroom and Ralston, 2016; Williams et al., 2019), we found fluorescent labeling of motor end plates was retained when fluorescent α-BTX was applied to the muscle prior to immersion fixation. Intrafusal endplates were clearly distinguishable from extrafusal neuromuscular junctions. γ-neuromuscular junctions are isolated and found in relatively close proximity to the equatorial annulospiral endings. On the other hand, α-neuromuscular junctions tend to be organised in grapes perpendicular to the main nerve branch (Figure 4A–C). Extrafusal endplates also tended to be much brighter and more complex than intrafusal endplates (Arbuthnott et al., 1992; Banks et al., 1985; Kucera and Walro, 1986). Based on these morphological differences, we were able to identify and study both α and γ neuromuscular junctions (Figure 4B–C). Optical clearing of the muscle also gives an unprecedented access to the three-dimensional structure of the annulospiral endings of the Ia afferent fibers around the equatorial region of the bag fibers. Z-stacks obtained on a confocal microscope highlight the complex organization of these structures (Figure 4D) and allow us to evaluate their morphology. While this study focuses on establishing the imaging workflow, the morphometric features that are now accessible using this approach are as follows. Discernible properties of the spindle equator included gross length and diameter, number of intrafusal fibers, which corresponds to the number of spirals, as well as Ia and II afferent diameters at spindle entries. Observable morphological features of spindle annulospiral endings included number of circumferential elements, their width, and their inter-rotational distances.

**Figure 4:**
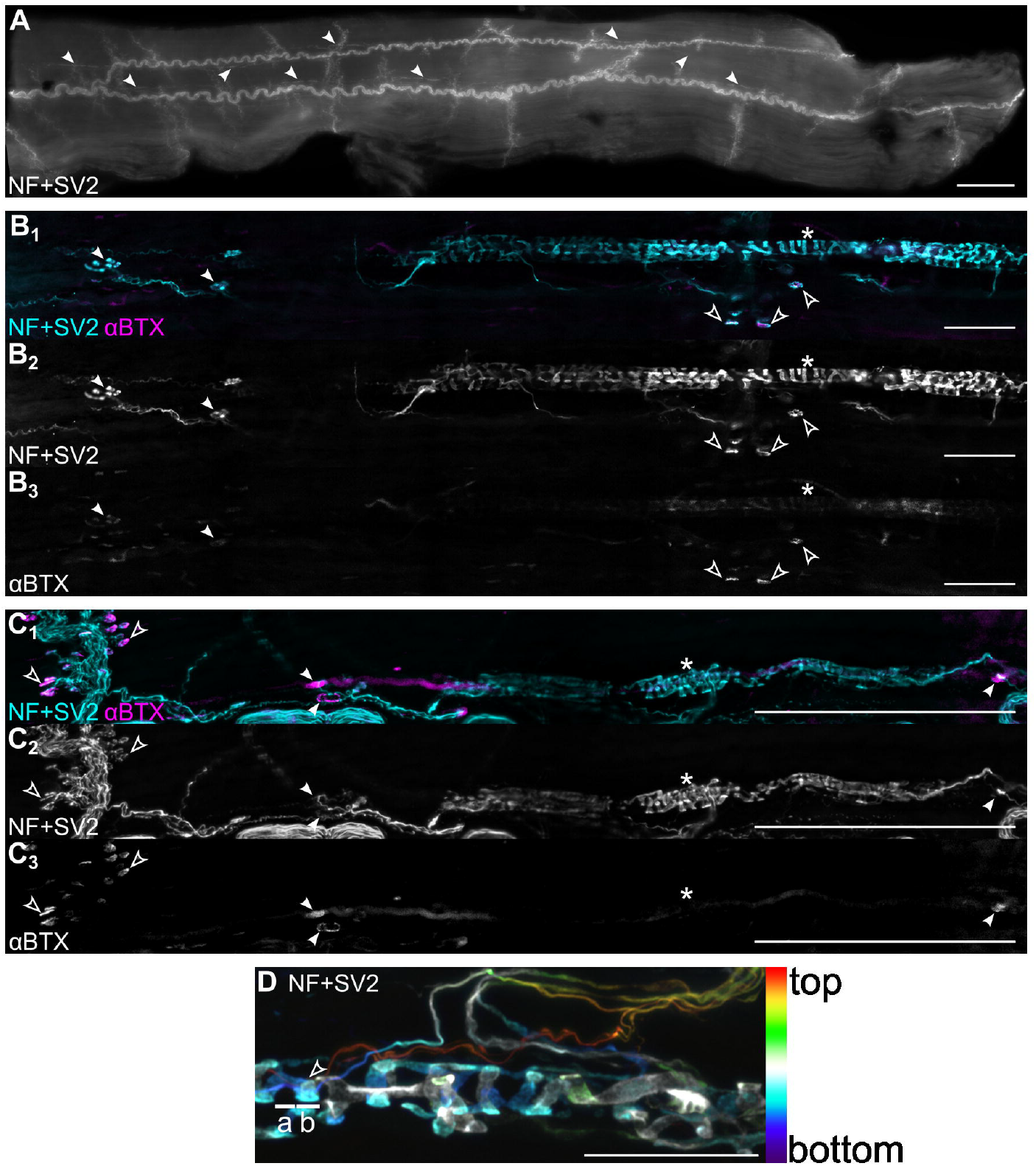
Representative images obtained from optically cleared tenuissimus muscles. **(A)** Maximal-intensity projection of a tiled (8×3 tiles) Z-stack (step size 20 μm, seven steps) epifluorescence image of a 8 day-old rabbit tenuissimus that was cut in half prior to optical clearing. In this half-muscle, at least nine spindles are readily visible (arrowheads), running parallel to the main nerve branch that serpentine through the whole length of the muscle. 10×/0.4 NA objective. Scale bar: 500 μm. **(B)** Confocal image of a single spindle from the same specimen as in (A) showing the overlay (**B**_**1**_) of the pre-synaptic element labeling (Neurfilament and SV2, **B**_**2**_) and the post-synaptic receptors (αBTX, **B**_**3**_). The asterisk * points to the equatorial region of the spindle. Empty arrowheads point to extrafusal neuromuscular junctions, while filled arrowheads point to putative γ endplates. Scale bar: 50 μm. Tiled image acquired at 40×/0.95NA (pixel size 6.1313 px/μm), pinhole size: 40 μm. **(C)** Confocal image of another spindle. Same organization as in B. Scale bar: 200 μm. Maximum-intensity projection of a tiled z-stack (1.25 μm step size, 16 steps) acquired at 20× (pixel size 3.0591 px/μm), pinhole size: 40 μm. **(D)** Depth-coded projection of a z-stack (0.6 μm step size, 33 steps) of the equatorial region of a spindle acquired with a 40×/0.95NA objective and a 1.5× optical zoom (pixel size 13.9844 px/μm). The color bar on the right shows how each color corresponds to a particular depth in the z-stack. Discernible morphology includes an example annulospiral circumferential element (arrowhead), inter-rotational distance (a), and coil width (b). Pinhole size: 40 μm. Scale bar: 50 μm.

Although confocal imaging provides the highest level of detail, it is limited by the working distance of the objectives, which is sometimes too short to focus on the spindles deep inside the muscle, particularly in older animals. Lightsheet microscopy was used to produce high resolution volumetric images of the muscle (Supplemental Movie 1), including muscle spindles (Supplemental Movies 2 and 3). The combination of optical clearing and lightsheet microscopy thus allows the study of the three-dimensional structure of muscle spindles in a wide range of ages, regardless of the thickness of the muscle.

## 4 Discussion

Several decades ago the tenuissimus muscle was the subject of foundational research on muscle spindle physiology and fusimotor innervation, due to its high spindle density (Banks, 1991; Barker et al., 1976; Boyd and Ward, 1975; Celichowski et al., 1994; Emonet-Dénand et al., 1975; Hunt and Kuffler, 1951, p. 1951). The tenuissimus generates very small forces because it is a thin muscle with few extrafusal fibers (Lev-Tov et al., 1988). It contains many spindles aligned in series near the midline of the muscle, parallel to the central neurovascular bundle (Figure 4A)(Miyoshi et al., 1979; Patten and Ovalle, 1992).This anatomical feature of the tenuissimus and its small size support the idea posed by (Patten and Ovalle, 1992) that the high degree of proprioceptive feedback provided by this muscle fine tunes gross movement in the neighboring biceps femoris muscle, which has a much larger force generative capacity (Patten and Ovalle, 1992; Peck et al., 1984). In this study, we adapted contemporary immunofluorescence and optical clearing methods for three-dimensional visualization of tenuissimus muscle spindles *in situ*. Our work returns attention to the tenuissimus and highlights its unique suitability for investigations in muscle spindle biology.

Muscle spindle morphology is dynamic. The sensory nerve terminal transitions from a web-like to a spiral appearance during early postnatal spindle development in rodents (Kröger and Watkins, 2021; Maeda et al., 1985; Osawa et al., 1988), and in aged mice, sensory axon blebbing, Ia afferent atrophy at muscle spindle entry, and reduced spirality of annulospiral endings has been reported (Kawai-Takaishi and Hosoyama, 2025; Kim et al., 2007). Morphometric analysis of muscle spindles typically involves labor-intensive teasing of silver-impregnated muscle spindles, longitudinal muscle cryosectioning, or serial transverse cryosectioning with three-dimensional reconstruction (Gartych et al., 2021; Gerwin et al., 2020; Kawai-Takaishi and Hosoyama, 2025; Kim et al., 2007; Muller et al., 2008; Piotr et al., 2023). Selecting muscles amenable to whole-mount microscopy, like the extensor digitorum longus (EDL) as done by (Vaughan et al., 2015), decreases labor intensity and maximizes the number of spindles for morphometric analysis. More recently, (Bornstein et al., 2022) demonstrated the great utility of optical clearing for *in situ* visualization of muscle spindles in the EDL, which has a reported spindle density of 1.80 spindles/gram of muscle in the adult mouse (Lian et al., 2022). In the current study, we adapted optical clearing for *in situ* spindle visualization in the tenuissimus, which has a very high spindle density (503 spindles/gram of muscle in the adult hamster)(Patten and Ovalle, 1992). The high spindle density of the tenuissimus makes it particularly well-suited for spindle morphological analysis.

Our workflow enables volume imaging of the entire spindle structure, including its fusimotor innervation (Figure 4). Others have used transgenic fluorescent reporter lines like *Thy1-eGFP* to visualize spindle annulospiral endings in mouse skeletal muscle optically cleared with hydrophilic-based (CUBIC (Shi et al., 2025) and SeeDB (Nikolaou et al., 2015)) and hydrogel-based (PACT (Bornstein et al., 2022)) methods, but none have labeled spindle poles (γ motor end plates) or used immunofluorescence to label spindle innervation. The protocol developed in this study may be useful for other groups studying spindle structure and/or those more broadly interested in visualizing neuromuscular junctions, particularly in species where no transgenic reporter lines exist.

Fluorescent dye-conjugated-α-BTX binds to the α subunit of nicotinic acetylcholine receptors and is the gold standard reagent for labeling motor end plates at α- and γ-neuromuscular junctions in skeletal muscle (Anderson and Cohen, 1974). While widely used in standard immunofluorescence experiments, fluorescent dye-conjugated α-BTX is challenging to pair with optical clearing of skeletal muscle. This reagent is incompatible with hydrogel-based optical clearing methods when the key component sodium dodecyl sulfate (SDS) is used for delipidation (Milgroom and Ralston, 2016; Williams et al., 2019); exclusion of SDS in the MYOCLEAR hydrogel-based method yields strong α-BTX labeling, but causes weaker antibody penetration and high blue-to-red range autofluorescence which limits co-labeling immunofluorescence experiments to those involving spectral unmixing of far red light (Williams et al., 2019). Fluorescent α-BTX has only been shown to label motor end plates in rodent skeletal muscle optically cleared using solvent-based protocols when administered by intravenous administration (Chen et al., 2016; Qi et al., 2019b; Yin et al., 2019), and it has never been paired with iDISCO (Renier et al., 2014)). The intravenous approach for motor end plate labeling is complex, since tail vein administration of fluorescent α-BTX requires a dosage (0.3 μg/g) above the LD50 for the mouse (0.113 μg/g) and for staining efficacy requires a 1-2 h delay under anesthesia before euthanasia and transcardial perfusion with paraformaldehyde. To our knowledge, this is the first demonstration that fluorescent α-BTX can be chemically fixed at nicotinic acetylcholine receptors by simple immersion fixation in paraformaldehyde and that this fluorescent labeling of motor end plates is compatible with iDISCO and solvent-based optical clearing. This approach may be useful for those interested in studying neuromuscular junction biology (γ and/or α) in optically cleared skeletal muscle.

## Limitations

The workflow described here was developed and validated using the tenuissimus muscle, which was selected because of its exceptionally high muscle spindle density and its thin, whole-mount amenable architecture. That said, the approach should be readily adaptable to other muscles, particularly rodent skeletal muscles that are comparable in size or smaller than the rabbit tenuissimus. As with most solvent-based optical clearing protocols, tissue shrinkage was observed and is unavoidable. However, because shrinkage is expected to occur consistently across samples processed using the same pipeline, it does not preclude relative comparisons between experimental groups when uniform processing conditions are maintained.

## Conclusion

In this study, we present a novel workflow for fluorescent labeling and volume imaging of muscle spindles in the neonatal rabbit tenuissimus, a whole mount-amenable muscle that is remarkably well-suited for spindle morphometry due to its high spindle density. Our novel approach enables in situ investigations into muscle spindle structure, including the annulospiral endings and intrafusal neuromuscular junctions in the contexts of health and disease.

## 6 Author Contributions

ER: Conceptualization, Data curation, Investigation, Methodology, Validation, Visualization, Writing – original draft, Writing – review & editing. BM: Investigation. OO: Investigation, Visualization. CK: Investigation. JG: Investigation. EF: Investigation. KQ: Funding acquisition, Supervision. MM: Conceptualization, Data curation, Investigation, Methodology, Validation, Visualization, Writing – original draft, Writing – review & editing.

## 8 Acknowledgments

The authors gratefully acknowledge Harvard’s university MicRoN (Microscopy Resources on the North Quad) Core (RRID:SCR_022209), the Harvard Center for Biological Imaging (RRID:SCR_018673), and the Rhode Island Institutional Development Award (IDeA) Network of Biomedical Research Excellence from the National Institute of General Medical Sciences of the National Institutes of Health under grant number P20GM103430 through the Centralized Research Core facility (RRID:SCR_017685) for infrastructure and support.

## 10 Figure legends

**Supplemental Movie 1**

Volumetric 3D reconstruction of part of a tenuissimus muscle from a P14 rabbit, imaged using a MegaSPIM lightsheet microscope (detection objective 22×/0.7NA). Fluorescence from the Neurofilament+SV2 immunolabeling.

**Supplemental Movie 2**

Volumetric 3D reconstruction of part of a tenuissimus muscle from a P14 rabbit, imaged using a Zeiss Lightsheet 7 microscope (0.5×/0.16NA detection objective, lightsheet thickness 5.17□μm). Fluorescence from the Neurofilament+SV2 Alexa Fluor 555 immunolabeling is shown in cyan, while fluorescence from α-BTX-Alexa Fluor 647 is shown in magenta. The movie shows a zoom on the equatorial region of a spindle.

**Supplemental Movie 3**

Volumetric 3D reconstruction of part of a tenuissimus muscle from a P14 rabbit, imaged using a Zeiss Lightsheet 7 microscope (20×/1.0NA detection objective, lightsheet thickness 3.84□μm). Fluorescence from the Neurofilament+SV2 Alexa Fluor 555 immunolabeling is shown in cyan, while fluorescence from α-BTX-Alexa Fluor 647 is shown in magenta. The equatorial region of a spindle is visible in the center of the volume, while extrafusal neuromuscular junctions organised in their typical grape pattern are visible in the lower right corner.

